# Delay of gratification in non-human animals: A review of inter-specific variation in performance

**DOI:** 10.1101/2020.05.05.078659

**Authors:** Irene Susini, Alexandra Safryghin, Friederike Hillemann, Claudia A.F. Wascher

## Abstract

The ability to regulate and withhold an immediate behaviour in pursuit of a more preferred or valuable, albeit delayed, outcome is regarded as an important cognitive ability enabling adaptive decision-making in both social and asocial contexts. Abilities to cope with a delay in gratification have been investigated in a range of species using a variety of experimental paradigms. The present study attempts a first systematic evaluation of available experimental data from non-human animals, which is an essential basis for quantifying biological and non-biological factors (*e.g.* socio-ecology *versus* experimental design) affecting performance in delay of gratification tasks. Data were sourced from 52 separate studies, and a comprehensive overview of the available literature on delay of gratification in non-human animals is presented. We present data from 21 species, spanning across eight taxonomic order, with 1–9 species tested per taxonomic group. We highlight variation in experimental paradigms used to study delay of gratification abilities in non-human animals, both with regard to reward type or experimental setup, and discuss the implications for comparative analyses. We conclude that, at present, cross-species comparisons of delay of gratification abilities are hindered by a lack of consistency in experimental designs and low number of species tested across taxonomic orders. We hope to stimulate research on a more diverse set of species, and that future studies consider potential social and ecological factors that cause intra-specific variation in test performances, that is repeatedly seen across species.

## Introduction

Animals, including humans, are frequently faced with decisions that affect what options or rewards become available in the future (‘intertemporal choice’, Stevens et al., 2011). For example, in cooperative interactions, individuals often invest time and energy although no immediate benefits are derived from reciprocation. In a foraging context, individuals may refrain from eating fruits upon first encounter in order to have riper fruits available in the future (Rosati 2017). In competitive contexts, if aggression is to be avoided, subordinate individuals must often wait in order to obtain access to resources monopolised by higher-ranking conspecifics (Bräuer et al. 2007). Intertemporal choices are also applicable to mate choice scenarios, wherein a female must decide whether to mate with an immediately available partner or whether to wait for a higher-quality mate (Sozou and Seymour 2003; Fawcett et al. 2012). Waiting, however, bears risks, as resources might become depleted, or an individual may not survive long enough for the reward to be harvested (Stevens and Stephens 2010; Hayden 2019).

The ability to delay gratification is one of the most challenging forms of behavioural inhibition, and enables adaptive decision-making in both social and asocial contexts (Beran 2015, 2018). Performance in delay of gratification tests is assumed to vary across individuals and to be linked to traits regulating motivational and control processes (Casey et al. 2011). In humans, such abilities are correlated with healthy dietary habits and financial wellbeing (Mischel 2014). Delay of gratification abilities during childhood have been suggested to predict positive outcomes later in life, *e.g.* in the context of social and academic competence (Mischel et al. 1988), resilience (Shoda et al. 1990), mental health (Tangney et al. 2004), and physical health, financial wealth, and low criminal activity (Moffitt et al. 2011). However, a recent replication of the famous Stanford marshmallow test - during which a child was offered a choice between one small but immediate reward or two small ‘future’ rewards if they waited for a set period of time - challenged the direct link between delay of gratification performance as a child and achievements later in life. While an association was found between the two components, effects were sensitive to the inclusion of control variables describing (i) family and (ii) parental-educational and economic background (Watts et al. 2018); but see Michaelson and Munakata, 2019).

Since the second half of the 20^th^ century, the ability to delay gratification has attracted considerable scientific interest from an array of disciplines, ranging from human economics (Frederick et al. 2002), psychology (Mischel et al. 1989), pharmacology (Evenden and Ryan 1996), and neuroscience (Casey et al. 2011), to animal cognition and behavioural ecology (Kacelnik and Bateson 1997; Miller et al. 2019). Measures of delay of gratification for non-human animals are often interpreted in a comparative context, *i.e.* some species are considered more or less able than others to delay gratification, *e.g.* (Miller et al. 2019). In non-human animals, the ability to delay gratification is typically assessed using established experimental paradigms: accumulation of rewards, exchange of rewards, hybrid delay task, and an intertemporal choice task that offers an immediate and a delayed option (for a detailed overview of tasks see Stevens 2017; Beran 2018; Miller et al. 2019). Successful experimental trials typically require two components: the selection of a delayed reward over an immediate option, *i.e.* ‘delay choice’, and the ability to sustain the decision even when the alternative, immediate reward is maintained within reach throughout the delay, *i.e.* ‘delay maintenance’ (Stevens and Stephens 2010). The ability to delay gratification is commonly evaluated as a measure of maximum delay endured, *e.g.* (Pelé et al. 2010; Dufour et al. 2012; Auersperg et al. 2013; Hillemann et al. 2014). In this particular experimental paradigm, namely the exchange task, only those focal subjects that successfully wait at least once in a given condition are further tested, while subjects that fail to wait are removed from the experiment. Given such a design, the results of these studies reflect only the behaviour of a limited number of individuals in a low number of trials, which may not be representative of larger populations or of the species in general (Dufour et al. 2012; Hillemann et. 2014). Another measure for the ability to delay gratification that is reported frequently in non-human animal studies is the mean number of trials during which an individual was successful (*i.e.* managed to wait the required time) within a given delay condition. However, other measures of waiting performance have been used.

A major concern associated with the comparison of inhibitory control abilities among studies or between species is that measures of such abilities are not always correlated when different experimental paradigms are used (Addessi et al. 2013; Brucks et al. 2017). A meta-analysis by Duckworth and Kern (2011) on inhibitory control in humans revealed that cognitive inhibition tasks and delay of gratification questionnaires yielded moderately correlated measures of self-control (Duckworth and Kern 2011). Potential challenges associated with the assessment of individual inhibitory control abilities and comparing performances across tasks remain relatively unexplored in the non-human animal literature (but see: Bray et al. 2014; Marshall-Pescini et al. 2015; Müller et al. 2016). Further, at present, no formal agreement exists with regard to which measures can be reliably used to quantify the ability to delay gratification – *e.g.* does a minimum delay have to be endured, how many individuals need to endure the delay, and in what percentage of trials? It is also unclear whether inter-individual variations in maximum delay tolerated reflects biologically meaningful differences in non-human animals. Considering the complexity of inhibitory control and its components, it remains questionable whether any single measure can be regarded as a comprehensive or reliable evaluation of an individual’s inhibition capacities (Brucks et al. 2017).

The present paper offers a systematic review of the available literature on experimental studies reporting delay of gratification performances in non-human animals. We aim to synthesise the available empirical evidence and to investigate whether waiting performance, *i.e.* mean percentage of successful waiting trials, can be compared across different species from a range of taxonomic groups. We further aim to describe how different experimental paradigms can affect waiting performance in different species. We outline gaps in the literature, which include the observations that (i) only a few species from a narrow taxonomic range have been tested so far, and (ii) that there is little standardisation of both experimental paradigms and reporting of results. We hope that our review stimulates future studies that address and close the gaps in the literature we identified here, to eventually enable systematic explorative analyses of biological (*e.g.* socio-ecology) and non-biological (*e.g.* experimental paradigm) factors affecting delay of gratification abilities.

## Methods

### Search protocol and criteria for inclusion

Literature searches were performed through the Web of Science research platform on August 22, 2019. An initial search for ‘delay of gratification’, ‘delayed gratification’, ‘self-control’, ‘impulse control’, ‘impulsivity’, ‘inhibitory control’, or ‘intertemporal choice’ as keywords yielded 35,268 abstracts. A further search in Scopus for the keywords ‘delay of gratification’, ‘delayed gratification’, ‘self-control’, ‘impulse control’, ‘impulsivity’, ‘inhibitory control’, or ‘intertemporal choice’ in abstract, title, keywords limit to biological sciences yielded a further 1,095 abstracts. In addition to the 36,363 records found through database searching, 20 studies were identified through alternative sources, such as a recent review by Miller et al. (2019). Abstracts were screened for inclusion and more than 99 % of studies were discarded due to the inclusion of humans as study subjects (36,298 studies). A further 33 studies were excluded from analysis owing to the relevant data being unavailable; only studies that presented mean percentages of successful trials, *i.e.* number of trials in which the focal individual waited out of the total number of trials, and which specified duration of the delay were selected. Further, research with forced learning trials was excluded from this study. A total of 52 papers were included in the present systematic review. See Figure 1 for a diagram of the search results and study selection process.

**Figure 1.**
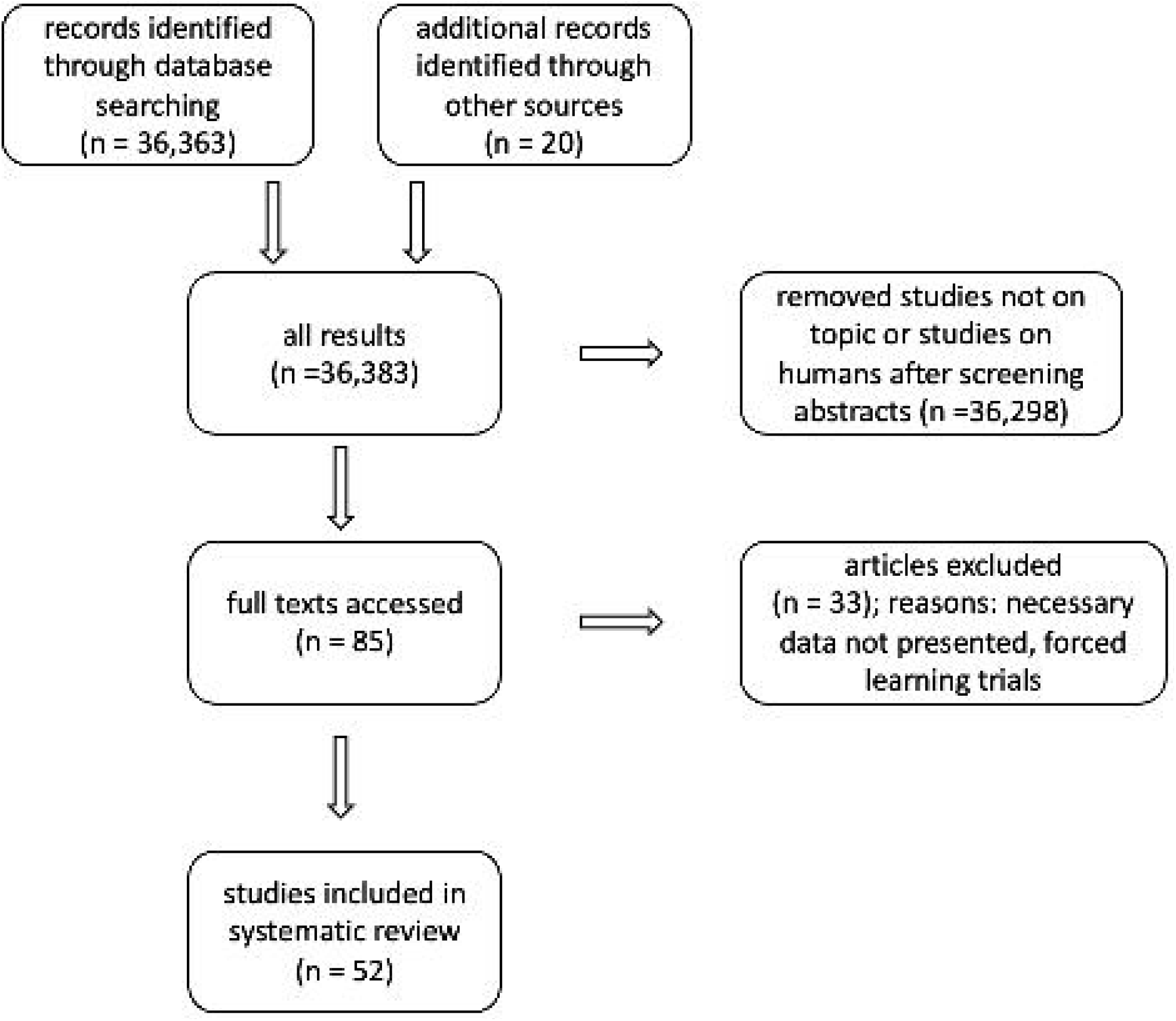
PRISMA diagram of the systematic literature and data search process.

From each included study, the mean percentage of successful trials of all tested individuals in a given delay condition was recorded. In the instance of focus animals being tested under multiple delay conditions, performances in each condition were recorded, resulting in multiple data entries per study. Experimental paradigm used (exchange, accumulation, go/no-go, intertemporal choice, rotating tray task) and reward type (qualitative, *i.e.* gain of a more preferred reward after a given delay; or quantitative, *i.e.* gain of more of the same rewards after a given delay) were also recorded. Mean percentage of successful trials was recorded for each focal individual in a given delay condition. The dataset generated and analysed in the current study are available as electronic supplementary materials.

### Descriptive Analysis

We describe the effect of a number of target factors on species’ performance in self-control tasks. Target factors included biological order, context (qualitative *versus* quantitative), amount of reward, duration of delay (seconds), form of delay test (maintenance *versus* choice), and experimental paradigm (accumulation, exchange, go/no-go, inter temporal choice, rotating tray); indicators of species’ performance included mean percentage of successful waiting trials and maximum endured delay. We refrained from a more formal explorative statistical analysis of the presented dataset due to the data being patchy and not balanced, as species were rarely tested in multiple different experimental paradigms.

## Results

Responses to delay of gratification tasks were analysed for 21 species - spanning across different taxonomic groups - tested under five different experimental paradigms. Primates are the most studied biological group, with nine species tested in total, followed by Passeriformes and Rodentia (five species each) and Psittaciformes (three species; Table 1). No species was tested in all five experimental paradigms; on average (± standard deviation, SD), species were tested in 1.4 (± 0.8) paradigms. The brown capuchin monkey, *Cebus apella*, was tested in four paradigms, making it the most-studied species. Only three of all tested species (14 %) were tested in both delay maintenance and delay choice tasks, and nine species (43 %) were tested in both quantitative and qualitative contexts.

**Table 1:**
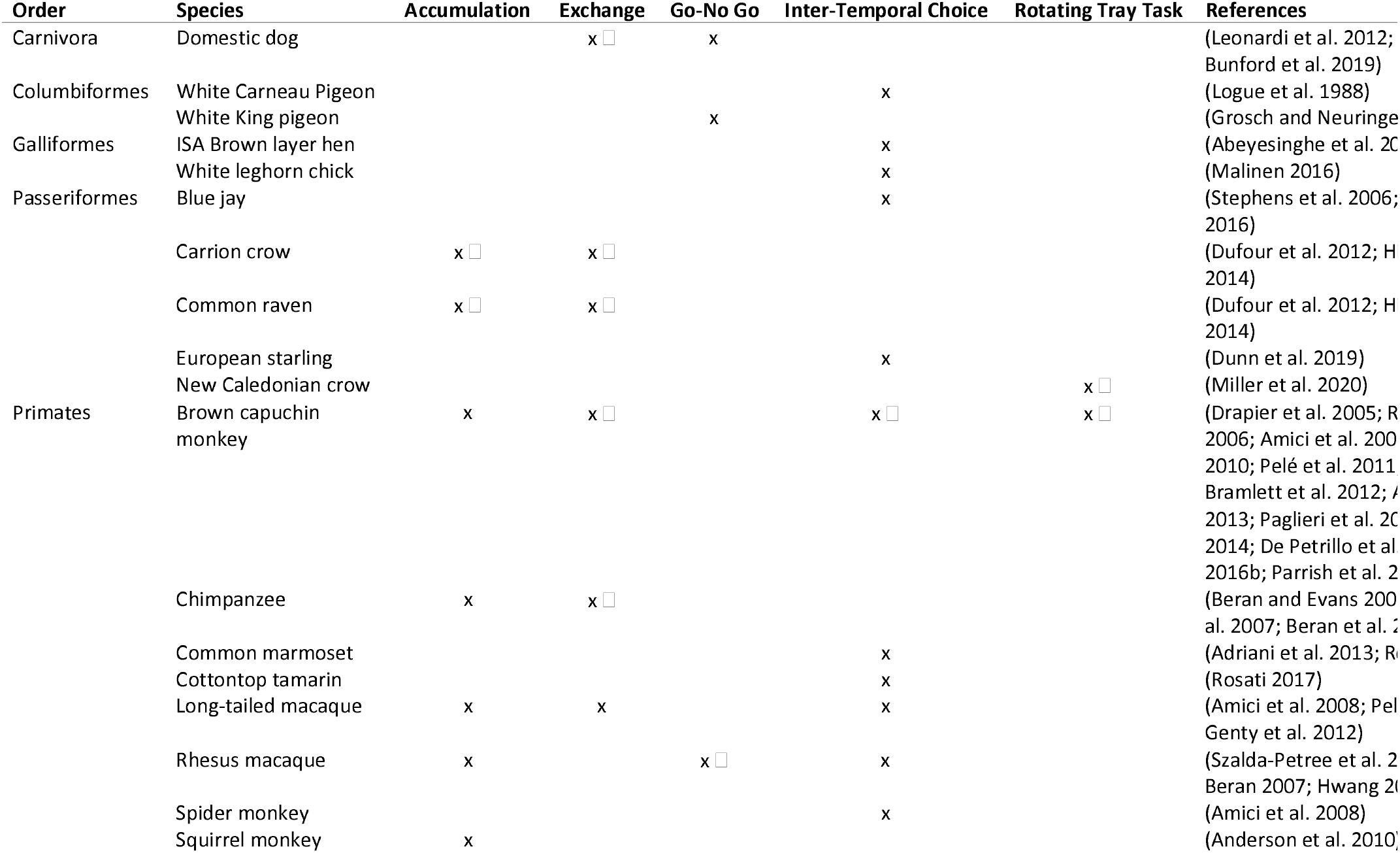
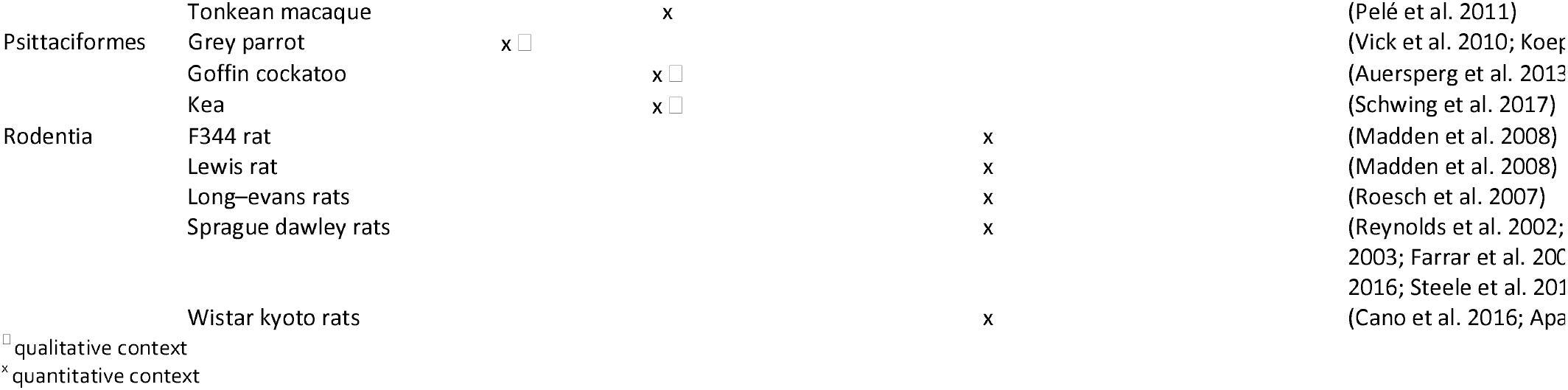
Different taxonomic orders and species tested in different experimental paradigms. X indicates a species has been tested in a specific paradigm, when the column is empty, the respective species has not been tested. Accumulation, exchange, go-no go and rotating tray tasks are delay maintenance tasks, and intertemporal choice tasks are delay choice.

Mean percentage of successful waiting trials ranged from 19 % successful trials in Passeriformes (*n* = 5) to 82 % successful trials in Psittacidae, Psittaciformes (*n* = 1). Range of performances in different orders and qualitative as well as quantitative tasks are shown in Figure 2. For most orders in which multiple species were tested under different paradigms, performance in tasks is highly variable (*e.g.* Primates, Passeriformes, Rodentia). Mean percentage of successful waiting trials decreased with duration of delay (Figure 3). Waiting performance varied in different experimental paradigms and was higher in delay choice (mean ± SD: 54 ± 23) compared to delay maintenance tasks (mean ± SD: 41 ± 33; Figure 4). Here, performance showed pronounced variation across different studies.

**Figure 2:**
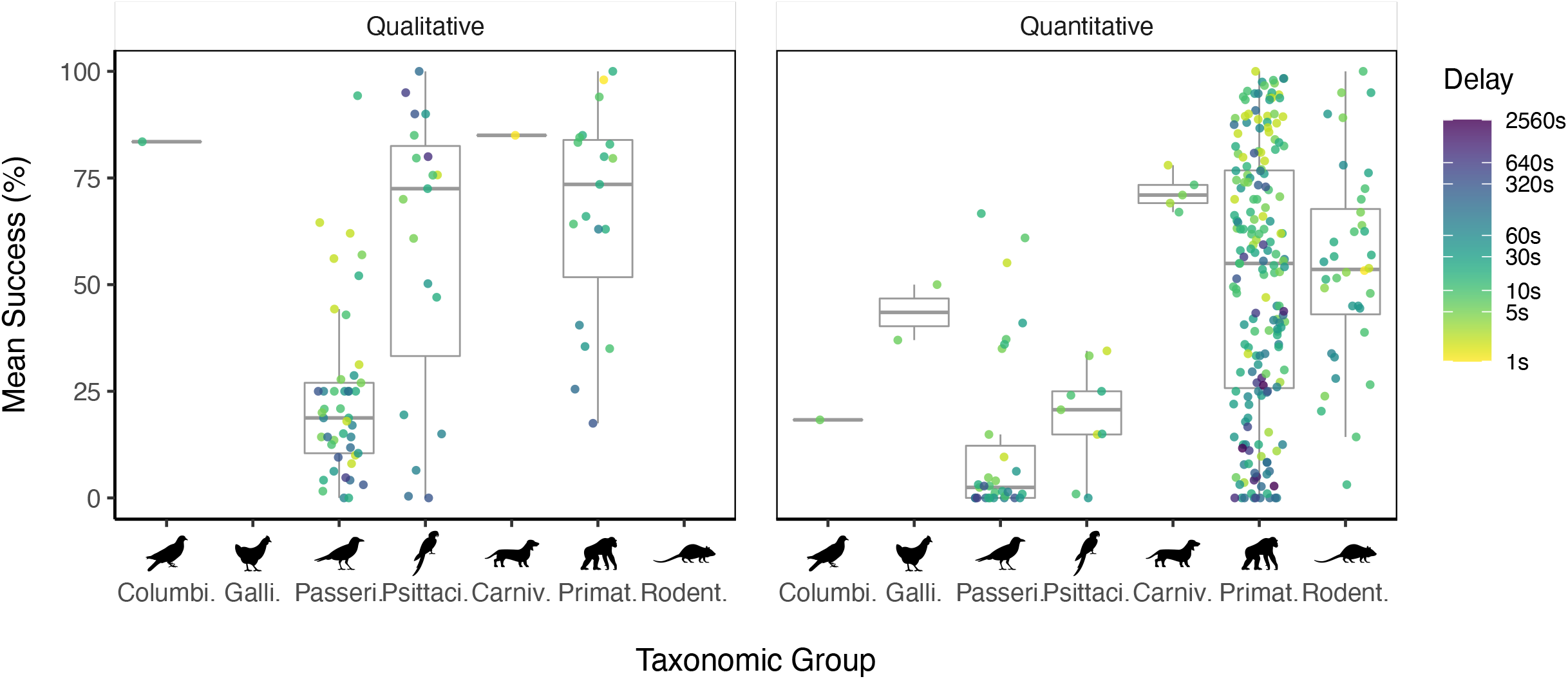
Mean percentage of trials in which individuals successfully waited in different biological orders. Box plots show the median of all individuals’ performances and the interquartile range from the 25th to the 75th percentiles, each data point represents the mean waiting performance per species per study in each delay condition. The colour of each data point codes the delay condition; note that the colour legend is not on a linear, but on a log scale. Across all species tested, individuals had higher rates of successful waiting in trials of shorter delays and performances dropped in longer delay conditions.

**Figure 3:**
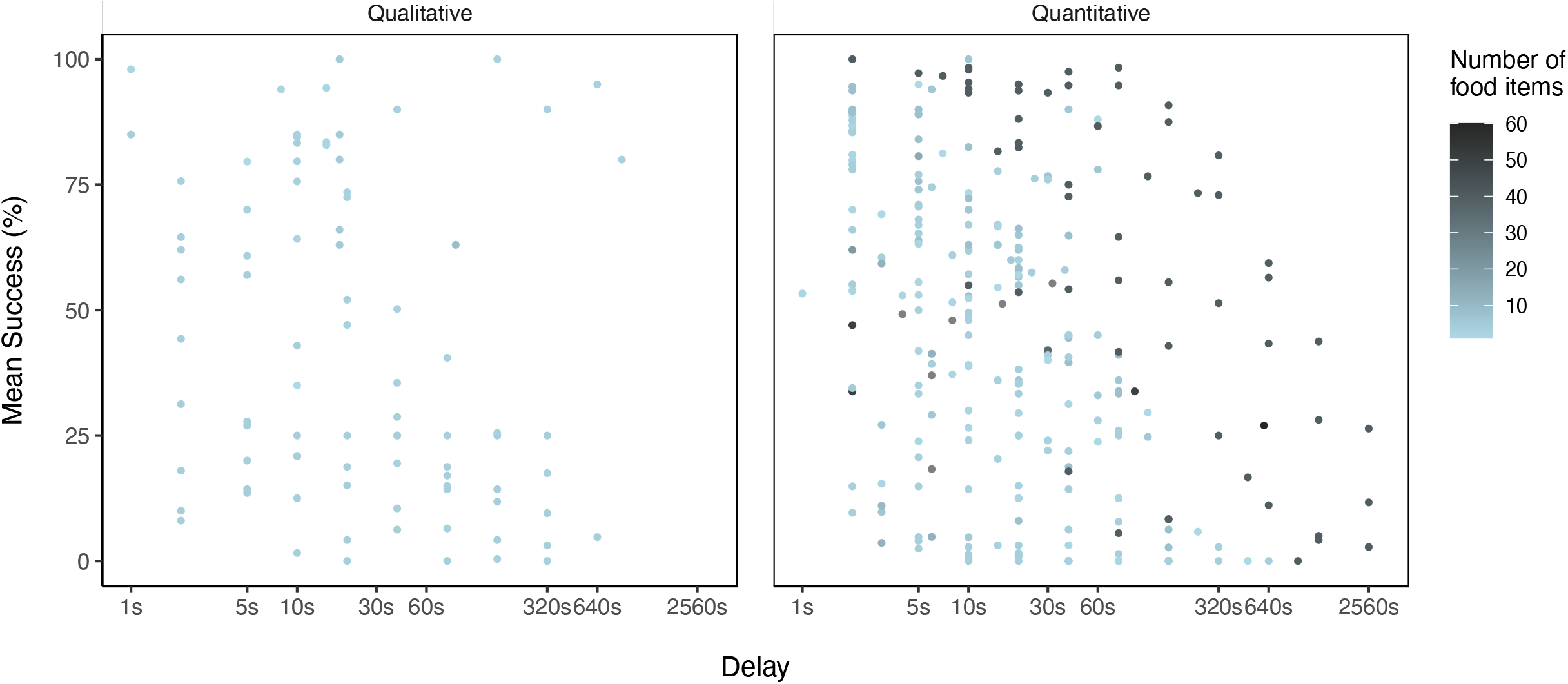
Mean percentage of trials successfully waited in a given delay condition. Data includes all species tested and individual dots present mean values per species per study. The shade of the dots represents the number of individual food items a subject gained in that experimental session. Note that the X-axis is on a log scale. With increasing delays, successful waiting performance decreases across all species, both when waiting for a qualitatively or quantitatively more preferred reward.

**Figure 4:**
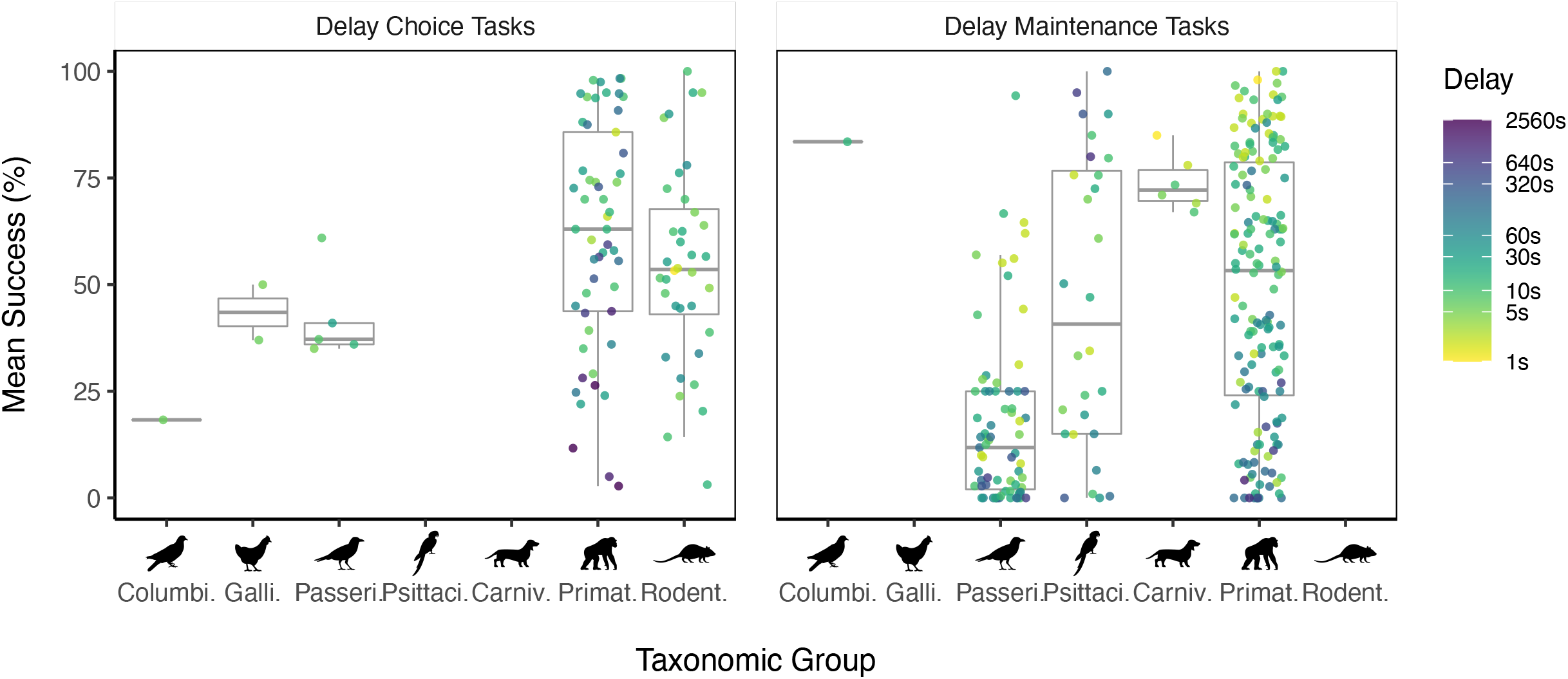
Mean percentage of trials successfully waited in different experimental paradigms. Box plots show the median and the interquartile range from the 25th to the 75th percentiles for each order (Columbiformes, Galliformes, Passeriformes, Psittaciformes, Carnivora, Primates, and Rodentia). The colour of each data point refers to the session’s delay condition; note that the colour legend is on a log scale.

## Discussion

The present study provides a systematic review of experimental data on delay of gratification abilities across different non-human species. It has been suggested that species differ in their abilities, or willingness, to wait for a better reward in food-related contexts due to socio-ecological factors. Here, we show that, due to a low number of individuals and species tested per taxonomic group, comparisons of biological factors affecting delay of gratification abilities should be taken with a grain of salt - if they are at all possible at present.

Delay of gratification studies typically report data for a small set of individuals (*e.g.* Beran and Evans 2006; Dufour et al. 2007, 2012; Pelé et al. 2010, 2011; Leonardi et al. 2012; Auersperg et al. 2013; Hillemann et al. 2014; Koepke et al. 2015; Schwing et al. 2017), which can bias estimates for a species or higher order, and may reduce the probability to detect between-species differences. The biological significance of inter-species differences in waiting performance remains, however, unclear. For example, Tonkean macaques, *Macaca tonkeana*, have proved capable of enduring delays of up to 2,560 seconds (Pelé et al. 2011), whilst carrion crows, *Corvus corone*, endured delays of up to 640 seconds (Hillemann et al. 2014). Does this mean that Tonkean macaques are four times better at coping with a delay in gratification compared to carrion crows, or that Tonkean macaques are able to delay future rewards whilst carrion crows are not? We suggest that mean delay performance, might be a more reliable measure of delay of an individual’s or a species’ gratification abilities compared to maximum delay endured; however, we also suggest that the development of more carefully standardised tests is necessary, and that these standardised tests should be conducted across a wider range of species. Our review of the available data suggests that previous conclusions about the evolution of cognitive skills under certain socio-ecological conditions are at best speculative, and that more comprehensive and systematic comparative studies are certainly desirable, as current results are drawn from a limited number of studies with a potential bias towards a small number of commonly tested species.

Our review reveals that the mean percentage of successful trials varies substantially across studies, depending on experimental design and on whether the design was a delay maintenance or delay choice task. The percentage of successful trials was higher in delay choice tasks than in delay maintenance tasks, suggesting that choosing to wait for a better reward is easier when the reward is removed after the focal individual has made its choice, compared to maintaining the wait while having access to a food item. Delay maintenance tasks often involve some sort of training procedure and are generally more time consuming, hence many researchers rely on delay choice tasks. However, delay maintenance represents a very interesting aspect of delay of gratification that includes many unaddressed questions. For example, do individuals tend to fail at the beginning or the end of the delay period, *i.e.* can they anticipate how long they will have to wait for, and can individuals develop distraction strategies that improve their test performance (Dufour et al. 2012).

A standardisation of experimental paradigms across taxa is challenging, as tasks differ in the degree of training and time required and are thus not universally applicable. For example, the exchange paradigms restricted to habituated and trained animals. Researchers have thus been calling for the development of new, automated procedures (Adriani et al. 2013; Miller et al. 2019). A further challenge associated with the implementation of standardised delay of gratification assessments is that artificial experimental designs often present alternative options simultaneously (*e.g.* smaller-sooner and larger-later), a setup which does not necessarily reflect food availability in natural foraging contexts and may encourage impulsivity (Pearson et al. 2010; Blanchard et al. 2013). Surprisingly, species which are viewed, generally, as likely to ‘fail’ to cope with delayed gratification under experimental conditions are regularly observed engaging successfully in natural behaviours that require such refraining abilities, including food caching, prey stalking, and long-distance travelling to high-quality food patches (Pearson et al. 2010). This raises the question of whether existing paradigms provide a reliable assessment of an animal’s delay of gratification abilities. Individuals’ performances in standard foraging problems (Hayden et al. 2011; Blanchard and Hayden 2014) may provide a more accurate measure of both human and non-human animals’ time preferences (Carter et al. 2015).

The present study provides a systematic overview of currently available data on delay of gratification performances in experimental studies and suggests that the data do not allow for conclusions to be formulated with regard to ecological and social factors driving the evolution of self-control. This is due to a lack of consistency in commonly utilised experimental paradigms, and a relatively small number of species tested. The implementation of experimental designs reflecting species-specific ecology is herein encouraged, in addition to the testing of more species that differ in their socio-ecological characteristics.

## Acknowledgements

We are grateful to Corina Logan and Dieter Lukas for feedback on an earlier version of the manuscript.

## Author contributions

Conceptualisation: IS and CAFW; Data curation: IS and CAFW; Formal analysis: FH and CAFW; Visualisation: FH; Writing: IS, AS, FH and CAFW.

## Notes

### Competing Interest Statement

The authors have declared no competing interest.

